# Natural variation in neuraminidase activity influences the evolutionary potential of the seasonal H1N1 lineage hemagglutinin

**DOI:** 10.1101/2024.03.18.585603

**Authors:** Tongyu Liu, William K. Reiser, Timothy J C Tan, Huibin Lv, Joel Rivera-Cardona, Kyle Heimburger, Nicholas C Wu, Christopher B. Brooke

## Abstract

The antigenic evolution of the influenza A virus hemagglutinin (HA) gene poses a major challenge for the development of vaccines capable of eliciting long-term protection. Prior efforts to understand the mechanisms that govern viral antigenic evolution mainly focus on HA in isolation, ignoring the fact that HA must act in concert with the viral neuraminidase (NA) during replication and spread. Numerous studies have demonstrated that the degree to which the receptor binding avidity of HA and receptor cleaving activity of NA are balanced with each other influences overall viral fitness. We recently showed that changes in NA activity can significantly alter the mutational fitness landscape of HA in the context of a lab-adapted virus strain. Here, we test whether natural variation in relative NA activity can influence the evolutionary potential of HA in the context of the seasonal H1N1 lineage (pdmH1N1) that has circulated in humans since the 2009 pandemic. We observed substantial variation in the relative activities of NA proteins encoded by a panel of H1N1 vaccine strains isolated between 2009 and 2019. We comprehensively assessed the effect of NA background on the HA mutational fitness landscape in the circulating pdmH1N1 lineage using deep mutational scanning and observed pronounced epistasis between NA and residues in or near the receptor binding site of HA. To determine whether NA variation could influence the antigenic evolution of HA, we performed neutralizing antibody selection experiments using a panel of monoclonal antibodies targeting different HA epitopes. We found that the specific antibody escape profiles of HA were highly contingent upon NA background. Overall, our results indicate that natural variation in NA activity plays a significant role in governing the evolutionary potential of HA in the currently circulating pdmH1N1 lineage.

## INTRODUCTION

Seasonal influenza A viruses (IAV) undergo rapid evolution in the human population, especially within the surface glycoproteins hemagglutinin (HA) and neuraminidase (NA)^1,2^. Substitutions within neutralizing antibody epitopes on HA, particularly in residues surrounding the receptor binding site (RBS), are thought to be the primary driver of viral antigenic evolution and the associated declines in vaccine efficacy^3–6^. Every 2-8 years, antigenically novel variants sweep across the globe, replacing previously circulating clades in a process known as an antigenic cluster transition^3,4,7^. These cluster transitions often coincide with elevated influenza-associated mortality rates^6,8,9^.

Given the high mutation rates of IAV observed *in vitro*^10–13^, it remains unclear why antigenic cluster transitions don’t occur more frequently. Numerous factors including narrow transmission bottlenecks, deleterious mutation load, and inter- and intra-segment epistatic effects likely all contribute to limiting the ability of the virus to explore novel antigenic space^14–18^. Identifying the specific factors that influence the emergence of antigenically novel variants is critical for improving the vaccine strain selection process and informing next-generation vaccine design.

During viral entry, HA forms multivalent bonds with glycoconjugates that contain terminal sialic acid residues^19–21^. These sialic acid receptors are common in the mucosal lumen, and NA plays an essential role in dissociating the virion from sialylated factors in the extracellular space or non-functional cell surface sialic acids to facilitate entry^22^. During virion release, NA liberates newly assembled virions from the sialic acids on the cell, preventing entrapment and aggregation, and promoting virus spread^23–25^. Thus, suboptimal HA-NA balance can affect the efficiency of both entry and release^22^. Numerous studies have suggested the importance of maintaining HA-NA functional balance during antigenic evolution, the evolution of drug resistance, and the emergence and adaptation of the 2009 H1N1 pandemic strain^26–30^.

We recently used the mouse-adapted H1N1 strain A/Puerto Rico/8/1934 (PR8) to demonstrate that the evolutionary potential of HA can be significantly constrained by epistatic interactions with NA^31^, suggesting that the need to maintain HA-NA functional balance may restrict antigenic variant emergence potential^22,32,33^. To assess the relevance of this phenomenon for recent seasonal H1N1 viruses, we generated recombinant viruses encoding the NA segments from H1N1 vaccine strains isolated between the pandemic lineage emergence in 2009 and 2019. We compared the NA activity phenotypes of these different N1 genotypes and asked how natural variation in NA activity altered both the mutational fitness landscape of HA1 by deep mutational scanning, and the genetic pathways of escape from neutralizing antibody selection. Our results demonstrate that epistatic interactions with NA can play a significant role in influencing HA antigenic evolution in the seasonal H1N1 lineage.

## RESULTS

### Variation in NA activity within the pdmH1N1 lineage

The pdmH1N1 NA gene accumulates ∼4×10^-3^ nucleotide substitutions per codon year, partially due to immune pressure against NA in the population^1,34–36^. To capture the evolution of the NA gene from its emergence into humans in 2009 to 2019, we selected five vaccine strains chosen by the WHO over this time period: A/California/07/2009 (CA09), A/Michigan/45/2015 (MI15), A/Brisbane/2/2018 (BR18), A/Hawaii/70/2019 (HI19), and A/Wisconsin/588/2019 (WI19) (**Fig. 1A**). 26 amino acid substitutions differentiate WI19 from CA09 for NA. We used reverse genetics to generate recombinant viruses that encode the NA genes from each of the five vaccine strains, with the remaining 7 segments coming from CA09.

**Figure 1:**
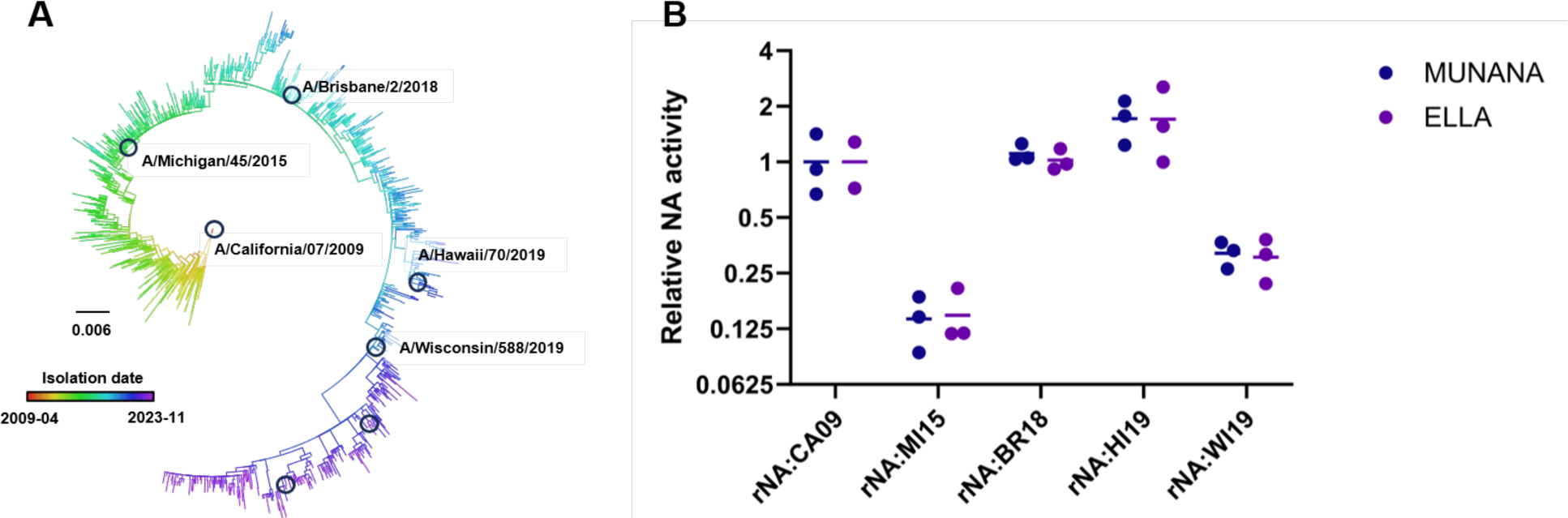
Relative NA activity varies across the pdmH1N1 lineage. **(A)** Phylogenic tree of the pdmH1N1 lineage NA segment with branch lengths in units of divergence (generated by Nextstrain^34^ and visualized in FigTree^37^). Vaccine strains marked with circles, strains used in this study are labeled with strain name. **(B)** Relative, virion-associated NA activities of recombinant viruses encoding the indicated NA genes in Cal09 backbone measured by either MUNANA or ELLA. Results were normalized based on genome equivalents of the NP segment as measured by qPCR.

We compared the NA activities of these viruses against the soluble NA substrate 2′-(4-Methylumbelliferyl)-α-D-N-acetylneuraminic acid (MUNANA), and against surface immobilized fetuin in the enzyme-linked lectin assay (ELLA) (**Fig. 1B**). We observed a nearly 10-fold range in virion-associated NA activity across the NA genes tested. Notably, the two vaccine strains isolated in 2019, HI19 and WI19, exhibited a roughly 4-fold difference in NA activity relative to each other, demonstrating how multiple NA genotypes with distinct NA activity phenotypes may co-circulate in a given year.

### Variation in NA background has little effect on the overall mutational fitness landscape of the pdm2009 lineage HA1

In our previous study, we observed that decreasing NA activity was associated with a global shift in the mutational robustness of HA, as measured by deep mutational scanning (DMS)^31^. To determine if the natural variation in NA activity we observed across the pdm2009 lineage could have the same effect, we used DMS to introduce all possible single amino acid substitutions in the HA1 subunit of the CA09 HA gene and compare the fitness effects of each substitution across different NA backgrounds.

We generated a hyper-mutagenized plasmid library containing all single possible amino acid substitutions in the HA1 subunit by degenerate PCR and rescued triplicate virus libraries for three different NA genes, CA09, HI19, and WI19, all in an identical CA09 backbone by reverse genetics (rNA:CA09, rNA:HI19, and rNA:WI19). We chose HI19 and WI19 as comparators because they (a) represent two NA clades (primarily associated with HA clades 5a.1 and 5a.2) that co-circulated from 2019 to 2022, and (b) exhibit a roughly 4-fold range in relative NA activity in our assays.

For comparison, we also generated triplicate rNA:CA09 virus libraries in the presence of the NA inhibitor zanamivir at a concentration we measured to inhibit 90% of NA activity (rNA:CA09 + NAI) (**Fig. S1**). All virus libraries were passaged once for 36 hours in MDCK-SIAT1 cells, starting at a low multiplicity of infection (MOI; 0.05 TCID50/cell). We collected viral supernatants and sequenced the post-passage virus populations along with the input plasmid library and wild-type plasmid by barcoded sub-amplicon sequencing^38,39^.

Out of 6846 possible mutations within the HA1 subunit, 6557 mutations had sufficient sequencing depth with good quality and were included in downstream analysis. We calculated relative fitness scores (representing relative growth advantages in our cell culture system) for each substitution based on the changes in frequencies between the plasmid library and the post-passage population. Given that silent mutations are likely neutral and nonsense mutations are likely lethal, we normalized relative fitness scores of all substitutions based on the mean relative fitness scores of all silent and nonsense mutations in each sample so that the mean silent mutation value was set at a score of 1 and the mean nonsense mutation value would have a score of 0. The normalized fitness scores of each mutation showed a high degree of correlation between replicate passages (Pearson correlations ranged from 0.72 to 0.81, **Fig. S2**), suggesting that our relative fitness effect estimates were reproducible across replicate populations.

The normalized relative fitness score distributions in the different NA backgrounds were bimodal, with a primary peak around a score of 1 and a smaller peak around the score of 0 (**Fig. 2**). In contrast with our previous results obtained using PR8^31^, the fitness score distributions were similar across the different NA backgrounds and the CA09+NAI control, suggesting that the range of NA activities tested here had little effect on the overall mutational robustness of HA1 in the pdm2009 lineage. The means of the average fitness scores from the different NA backgrounds were statistically significant (p < 0.001 by the Wilcoxon matched-pairs signed rank test); however, so were the differences between technical replicates for the individual groups, suggesting that these differences primarily reflect technical noise. Altogether, these data indicate that the NA genotype has minimal effect on the global mutational robustness of HA1.

**Figure 2:**
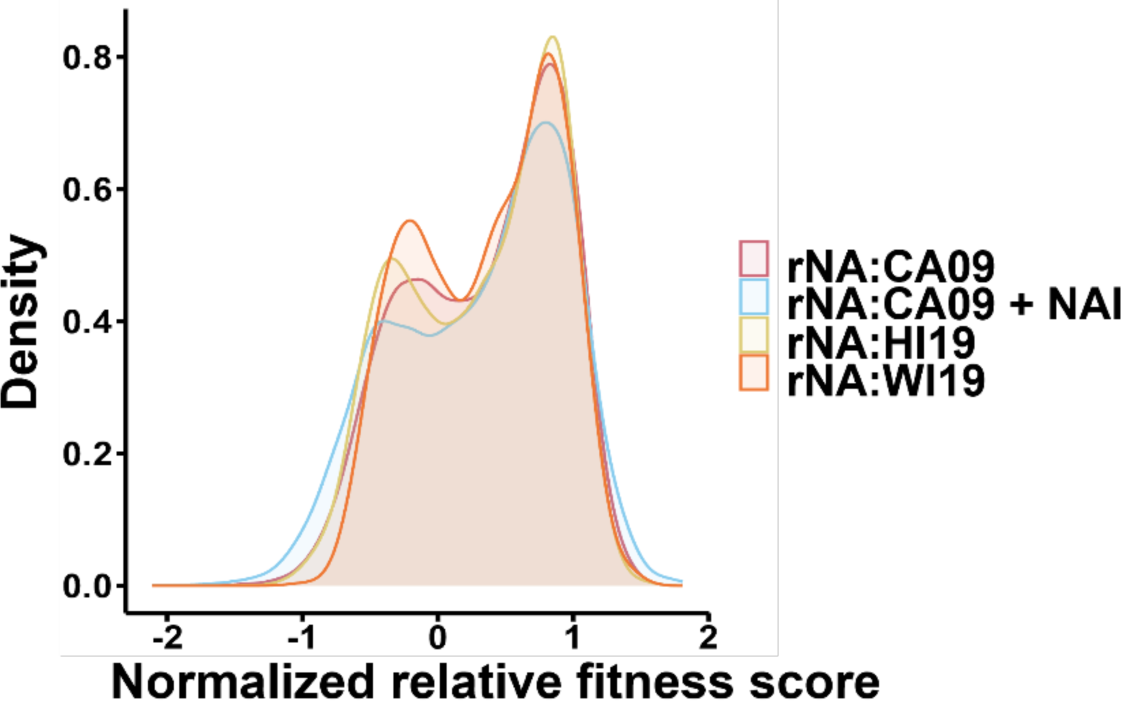
Changes in NA background have minimal impact on the overall mutational fitness landscape of HA1. The normalized relative fitness distribution of all missense mutations within HA1 subunit for each virus genotype tested, as well as the CA09+NAI control.

### Residues involved in receptor binding exhibit epistasis with NA

To determine whether changes in NA activity were associated with epistatic effects on specific HA residues, we performed linear regressions using the fitness score of each substitution as the dependent variable and the relative NA activity as the independent variable for the different NA backgrounds. Substitutions that exhibited higher relative fitness in the context of higher NA activity had positive coefficient values, and those with higher relative fitness in the context of lower NA activity had negative coefficient values. The absolute value of the coefficient quantifies the sensitivity of a given substitution to the epistatic influence of NA.

Given that both sialic acids and the exterior of the plasma membrane are negatively charged, we predicted that substitutions that increased positive charge on the HA surface would enhance attachment to the cell surface and sialic acids and thus would exhibit higher relative fitness in the context of higher NA activity. Similarly, we predicted that negative charge changes would be better tolerated in reduced NA activity backgrounds. Consistent with this, we observed significantly more positive charge changes with positive coefficient values than negative charge changes (**Fig. 3A**).

**Figure 3:**
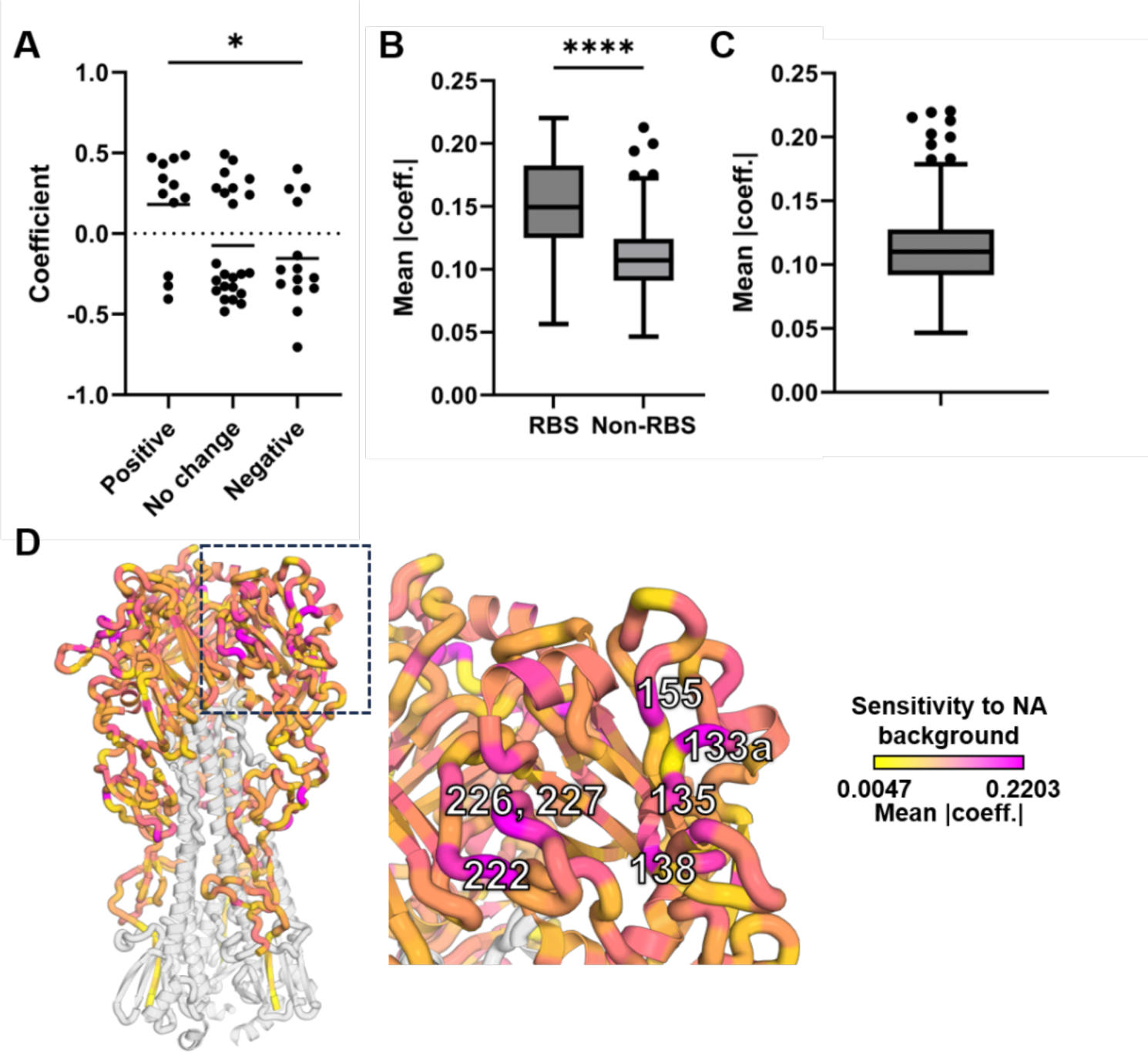
Epistatic interactions with NA are most pronounced in RBS-associated residues. **(A)** The coefficient values (p < 0.001) of different charge changes on the HA surface, * indicates p < 0.05 generated by Mann Whitney test. Substitutions with p>0.001 are not shown. **(B)** Comparison of mean of absolute coefficient (MAC) values of RBS-associated residues (defined as residues in the 130-loop, 190-helix, 220-loop or receptor binding pocket) versus the other residues in HA1, **** indicates p < 0.0001 generated by Mann Whitney test. **(C)** The Tukey box and whisker plot of MAC for all residues, outliers are shown as dots that are higher than 75th percentile plus 1.5 times IQR. **(D)** Structure of HA (PDB: 3UBQ^40^) colored by MAC value. The residue numbers on the RBS that were outlier high MAC values identified in (C) are labeled.

To quantify the overall epistatic influence of NA over each HA1 residue, we calculated the mean of the absolute coefficient (MAC) values across all nineteen possible substitutions for each residue. We hypothesized that substitutions at RBS-associated residues are more likely to have pronounced effects on receptor binding avidity than substitutions distal from the RBS and thus should be subject to stronger epistatic interactions with NA. We observed that the RBS residues exhibited higher MAC values than non-RBS residues (**Fig. 3B**). When we examined the overall distribution of MAC values across HA1, we identified residues 49, 133a, 135, 138, 155, 222, 226, 227 and 291 (all H3 numbering) as outliers, indicating that they are particularly sensitive to NA phenotype (**Fig. 3C**). With the exceptions of residues 49 and 291, all of these NA-sensitive residues are in or directly adjacent to the RBS (**Fig. 3D**), demonstrating that NA epistatic effects are concentrated in and around the RBS.

### Specific mutational pathways of HA escape from neutralizing antibody selection are contingent upon NA genotype

We next asked whether different NA backgrounds supported the emergence of distinct HA escape variants in the presence of identical neutralizing anti-HA monoclonal antibodies (mAbs), as we recently observed for the lab-adapted PR8 strain^31^. We used a panel of mAbs targeting distinct epitopes on HA (**Table 1**).

**Table 1.** Monoclonal antibodies used in the selection experiments.

| Antibody | Epitope | Reference |
| --- | --- | --- |
| EM4-C04 | Sa | 28,41 |
| 2A05 | Ca1 | 42 |
| 2C05 | Ca2 | 42 |
| CR9114 | Stem | 43 |

For each mAb, we determined the saturating neutralization concentration, defined as the lowest mAb concentration that maximized the inhibition of virus yield under the same experimental conditions (virus population size, volume, etc.) as the subsequent selection experiments **(Fig. S3)**. For each mAb, we incubated 10^7^ TCID50 of each recombinant virus (pooled from three independent virus rescue transfections) with each mAb at its saturating neutralization concentration in six replicates. We used starting populations of 10^7^ TCID50 to maximize representation of all possible single nucleotide substitutions in starting populations. This was based on the assumption that each replicated genome contains an average of 2 random nucleotide substitutions^10^, meaning that every possible single nucleotide mutation would occur ∼370 times on average within a starting viral population of 10^7^ TCID50 (ignoring the effects of variation in relative fitness and substitution type preferences of the viral replicase^11^). We infected cells with virus/antibody mixtures and collected virus supernatants at 24 hours post infection for next-generation sequencing to identify putative escape variants. If viral titers were insufficient (< 10^4^ TCID50/mL) for generating high quality sequencing libraries, we performed a second passage in the presence of the mAbs.

Based on our DMS results, we hypothesized that RBS-proximal epitopes would be more sensitive to changes in NA background. For the Sa-specific mAb EM4-C04^41,44^, K133aT was the dominant HA substitution that emerged in the rNA:CA09 background, while K133aI and K133aN were the major substitutions that emerged in rNA:HI19 and rNA:WI19 (**Fig. 4A, B**). Note that we identified K133a as one of the NA sensitive residues by DMS (**Fig. 3**). K133aN first began to emerge during the 2018-2019 flu season and swept to global dominance by 2023, raising the possibility that the emergence of this antigenic variant at the global scale may have depended in part on the emergence of suitable NA genotypes **(Fig. S4)**.

**Figure 4:**
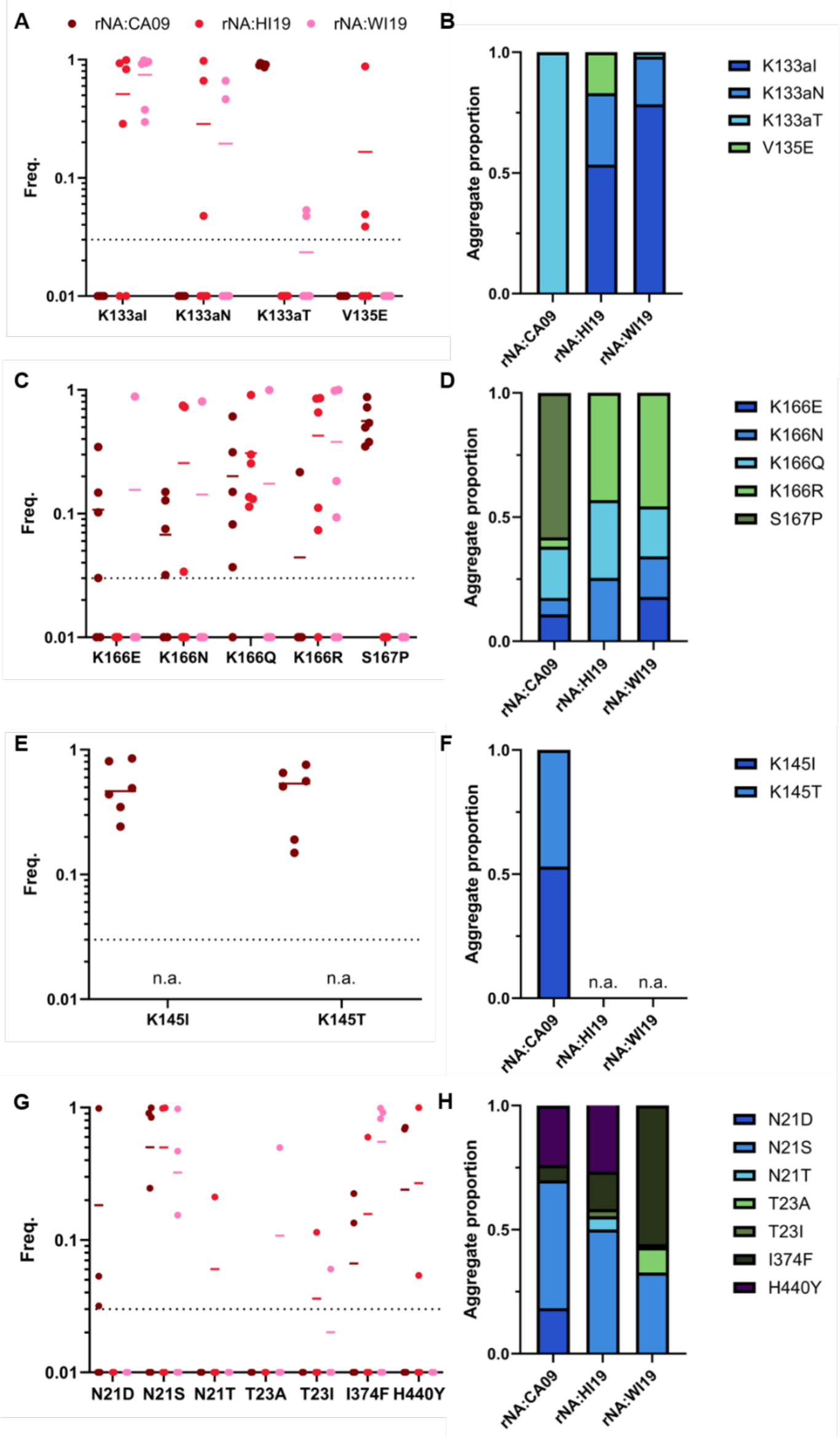
Specific mutational pathways of anti-HA antibody escape are contingent NA activity. HA substitutions that emerged under mAb selection in different NA backgrounds. **(A, C, E, G)** Frequencies of major substitutions (>3% in a given replicate and >10% in at least one replicate) detected under selection with (A) EM4-C04, (C) 2A05 (E) 2C05 (n. a., not assayed due to no surviving virus.) and (G)CR9114, in colored by NA backgrounds (each dot represents the frequency in a single selection replicate n = 6 replicates, except for CR9114 with rNA:WI19 (n = 4), 2A05 and CR9114 with rNA:WI19 (n = 5)). (B, D, F, H) The aggregate proportions of each indicated substitution summed across all replicates for each NA background.

Under selection with the Ca1-specific mAb 2A05, S167P was the major substitution found in rNA:CA09 and K166R was the major substitution found in the other two NA backgrounds **(Fig. 4C, D)**. Note that the escape profiles of rNA:HI19 and rNA:WI19 for both EM4-C04 and 2A05 are more similar to each other than to rNA:CA09, despite exhibiting a greater difference in virion-associated NA activity **(Fig. 1B)**. This suggests that our NA activity assays may not capture all features of NA relevant for influencing HA fitness in our system.

To compare with escape from head-targeted mAbs, we performed selections using the stem-targeting mAb CR9114^43^. We observed the emergence of substitutions at N21, T23, I374 and H440, all located in or under the HA stem epitope (**Fig. 4G, H**). N21S emerged to >80% in one or more replicates for all three NA backgrounds examined. I374F (I45F, HA2 numbering) also repeatedly emerged to high frequency (>10%) in one or more replicates across all NA backgrounds, suggesting that some anti-stem mAb escape pathways may be less constrained by epistatic interactions with NA than what we observe with RBS-proximal residues. We did observe some stem escape variants that differed between NA backgrounds, including N21D, which was observed in multiple rNA:CA09 replicates, but not with rNA:HI19 or rNA:WI19, and H440Y (H111Y, HA2 numbering), which emerged in multiple rNA:CA09 and rNA:HI19 replicates, but not rNA:WI19. Interestingly, H440Y is not surface exposed - it is located underneath the stem epitope, adjacent to I374. This suggests that substitutions at buried residues may trigger conformational shifts within the stem epitope sufficient to mediate escape.

Finally, under selection with the Ca2-specific mAb 2C05, K145I and K145T both emerged in the rNA:CA09 background; however, no virus could be recovered post-selection from the rNA:HI19 and rNA:WI19 backgrounds **(Fig. 4E, F)**. This suggests that suboptimal NA pairings may limit the potential for evolutionary rescue in the face of some anti-HA immune pressures. When we compared post-selection titers between NA backgrounds, we observed substantial differences depending on the mAb used. No mAb control titers for rNA:HI19 and rNA:WI19 were about 10-fold lower than that of rNA:CA09, indicating that rNA:HI19 and rNA:WI19 have a slight intrinsic growth disadvantage compared to rNA:CA09 in the absence of Ab pressure **(Fig. 5)**. This can’t fully explain the much larger differences in post-selection titers for EM4-C04 and 2C05. This appears to be a mAb-specific phenomenon, as post-selection titer differences for 2A05 and CR9114 were comparable to no Ab controls.

**Figure 5:**
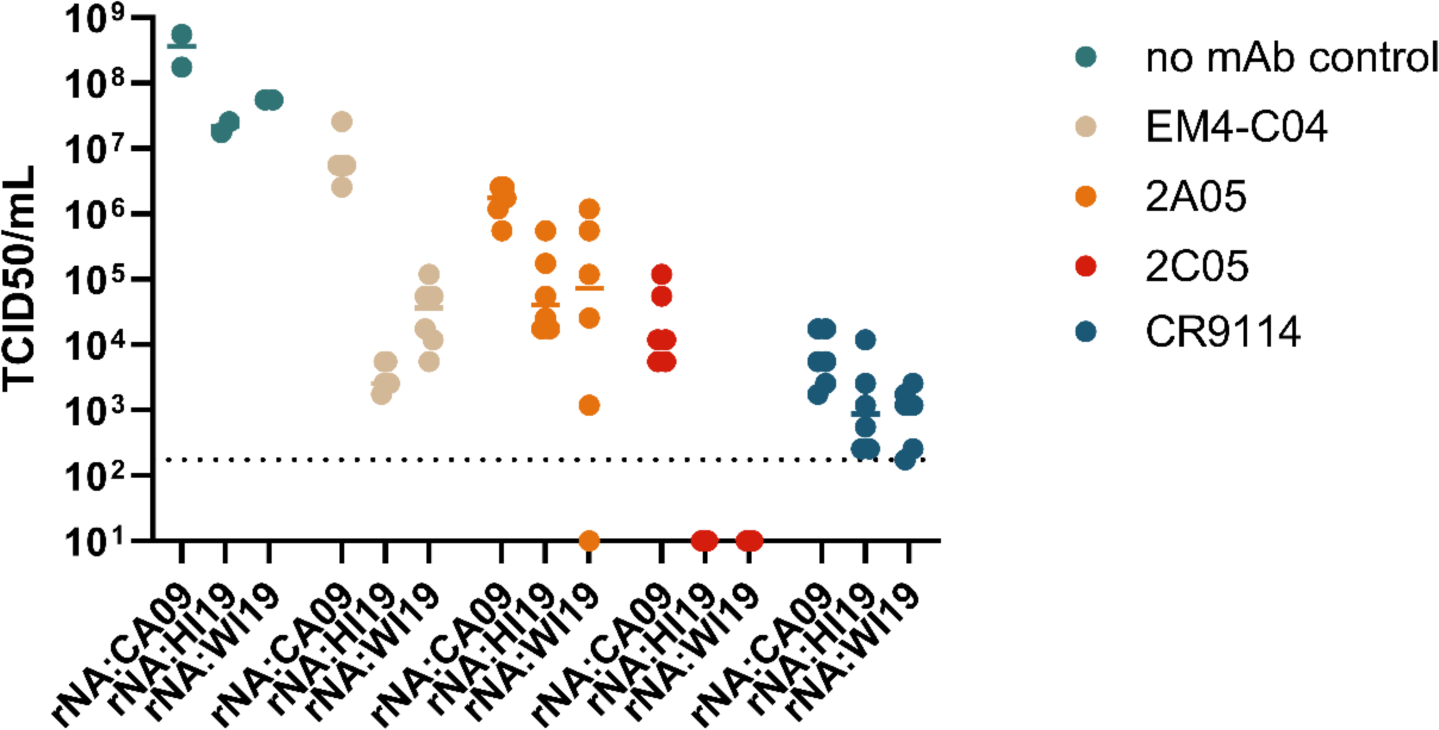
Differences in post antibody selection titers depending on NA background. TCID50 titers of viruses with different NA backgrounds (n = 6 for selection, n = 2 for no antibody control) following monoclonal antibody selection.

The specific dominant escape variants that emerged in different NA backgrounds under Ab selection could not be predicted based on the normalized relative fitness scores from the DMS experiment **(Fig. S5)**. For instance, S167P was the dominant variant that emerged in the rNA:CA09 background under 2A05 selection, despite having a lower relative fitness score than other possible escape variants. This discrepancy between the DMS results (which simply reflect relative growth advantages of individual substitutions) and the Ab escape results (which reflect a combination of mutation rates, initial population frequencies, growth advantage, and magnitude of Ab escape phenotype) illustrate how DMS-based approaches alone are insufficient for predicting the evolutionary trajectories of viral populations.

## DISCUSSION

Our results here extend our earlier proof-of-principle experiments with the lab-adapted PR8 strain^31^ to demonstrate that natural variation in NA within the pdm2009 H1N1 lineage can influence the specific genetic pathways taken by the virus to escape from anti-HA antibody pressure. These data clearly establish how the need to optimally tune the intimate functional partnership between HA and NA imposes distinct constraints the evolutionary potential of recent seasonal H1N1 viruses.

When we examined the distribution of normalized mutational fitness effects within HA1 using DMS, we observed that the major peak was centered around a score of 1, consistent with a large fraction of potential substitutions being nearly neutral in our system. That finding contrasts with WT PR8, where we observed the primary peak close to 0, indicating that most substitutions in that genetic background were highly deleterious or lethal^31^. This suggests that the pdm2009 H1N1 lineage HA1 is more mutationally tolerant than that of PR8. Given that the degree of mutational robustness of a gene can influence its potential to access fitness peaks^45^, more work should be done to quantitatively compare mutational fitness effect distributions between different virus strains and across different host environments. Significant differences in these distributions could suggest that different viral strains vary in overall evolutionary potential.

Also contrasting with PR8, we did not observe any significant shifts in the fitness effect distribution between different NA backgrounds (**Fig. 2A**). While we failed to detect a clear epistatic influence of NA over the global HA1 fitness landscape, we did detect strong epistatic effects for specific residues, primarily in or around the RBS. This is particularly important since RBS-proximal substitutions play a large role in driving antigenic cluster transitions3. This observation, combined with our observation that the pdm2009 HA takes distinct mutational pathways to escape from neutralizing antibody pressure depending on the NA background, strongly suggests that the antigenic drift potential of seasonal H1N1 viruses is significantly influenced by epistatic interactions between HA and NA.

While our DMS analyses clearly identified residues with mutational fitness effects that correlated with NA activity, our Ab escape experiments did not reveal a simple relationship between measured NA activity and escape variant emergence. For the head-specific mAbs EM4-C04, 2A05, and 2C05, we observed that rNA:HI19 and rNA:WI19 supported the emergence of more similar escape variants profiles to each other than either did to rNA:CA09. This does not line up with our NA activity data, where rNA:CA09 exhibited an intermediate phenotype in between those of rNA:HI19 and rNA:WI19. These data suggest that while NA can exert profound epistatic effects on the evolution of Ab resistance by HA, this effect cannot be simply explained by NA activity (at least as measured by the standard *in vitro* assays we use here). Future efforts should explore whether other NA features (*e.g.* receptor specificity, physical arrangements on virions, etc.) influence HA-NA epistasis.

We initially hypothesized that HA escape from stem Abs would not be significantly influenced by NA due to the distance between the stem epitope and the RBS. While the escape profiles for the three NA backgrounds did share some variants in common, namely N21S and I374F, rNA:WI19 differed from rNA:CA09 and rNA:HI19 both in the emergence of I374F at much higher frequencies, and in the absence of H440Y. These results suggest that NA may influence antibody escape at the stem epitope, despite its distance from the RBS.

We note that H440Y emerged under selection with CR9114 in both rNA:CA09 and rNA:HI19 backgrounds, despite being surface buried. H440Y is adjacent to the surface residue I374 where substitutions can convey resistance to CR9114 neutralization (as shown here and in a previous study^46^). Mutations at this position also promote the escape of H5 from CR6261^47^, a stem antibody encoded by the same germline gene GHV1-69 as CR9114. These data demonstrate how substitutions at residues buried underneath the stem epitope can mediate anti-stem Ab escape by inducing allosteric changes in the local structure that disrupt Ab binding.

Altogether our data highlight how epistatic interactions with NA constrain the evolutionary potential of HA in the seasonal pdm2009 H1N1 lineage. These findings indicate that a better understanding of HA-NA functional interactions in seasonal IAVs may help improve our ability to predict future evolutionary dynamics and enhance the accuracy of vaccine strain selection decisions. In practical terms, this will require more comprehensive genetic and phenotypic analysis of co-circulating HA and NA genotypes to determine whether HA-NA functional balance phenotypes can be inferred from sequence data and whether these phenotypes are predictive of the evolutionary success of specific sub-lineages.

## MATERIALS AND METHODS

### Cells and virus

HEK-293T and MDCK-SIAT1 cells were carried in Minimum Essential Medium (MEM + GlutaMAX, ThermoFisher Scientific) with 9.1% fetal bovine serum (FBS, Avantor Seradigm Premium Grade Fetal Bovine Serum) in 37 °C and 5% CO_2_.

Expi293F cells (Gibco) were grown and maintained in Expi293 Expression Medium (Gibco) at 37 °C, 5% CO_2_ and 95% humidity with shaking at 125 rpm.

Virus was first rescued by reverse genetics. 500 ng of each of 8 plasmids that transcribe both positive- and negative-sense viral RNA were used to transfect HEK-293T cells in 6-well plate by jetPRIME (Polyplus Transfection) according to the manufacturer’s protocol. 24 hours post transfection, the medium was replaced by infectious medium (MEM + 1 μg/mL TPCK-treated trypsin + 1 mM HEPES and 50 μg/mL gentamicin). MDCK-SIAT cells were co-seeded if necessary. The supernatant was then collected 48 hours post transfection and used to infect confluent MDCK-SIAT cells in 6-well plates. For infection, MDCK-SIAT cells were first washed once with PBS and the virus supernatant was added on top of the monolayer. After 1 hour of inoculation in 37 °C on shaker, the monolayer was washed again with PBS and infectious medium was added on the monolayer. The supernatant was collected 24-48 hours post infection as the seed stock of the virus. To generate the working stock of the virus, confluent T75 or T175 flask of MDCK-SIAT1 were infected with MOI of 0.0001 based on the TCID50 titer to avoid the generation of defective interference particle, and supernatant was collected 48-72 hours post infection.

One adaptive mutation to our cell culture system on HA (HA:G158E) was constantly found in high frequency in the working stock and readily fixed after few passages. We decided to introduce this mutation into the virus as the new CA09 backbone that would otherwise unavoidably emerge in our experiment. The numbering of HA and NA is based on the alignment of H3N2 amino acid sequence (strain A/Hong Kong/1/1968 H3N2, UniProt: Q91MA7, Q91MA2 for HA and NA) without the localization signal peptide unless specified otherwise.

### TCID50 assay

96-well plate was seeded a day prior to the experiment with 4-5×10^4^ MDCK-SIAT1 cells per well. In the day of the experiment, cells were washed with PBS once and 180 μL of infectious medium was added into each well. 20 μL virus was added into the first four rows of first column and continue with 10-fold serial dilution. Between each dilution, tips were changed to avoid carrying-over. The plates were scored for the endpoint of CPE of each row 5-7 days post infection. The TCID50 titer were calculated by Reed and Muench method^48^.

### Quantification of virus particle concentration by RT-qPCR

To remove extra-cellular viral RNA, 140 μL virus supernatant was treated with 0.25 μg RNaseA for 30 min in 37 °C and continued with viral RNA extraction (QIAamp Viral RNA Kits, Qiagen) following the manufacturer’s protocol. cDNA of the virus genome was generated by reverse transcription (Verso cDNA Synthesis Kit, ThermoFisher Scientific) using MBTuni-12 (5′-ACGCGTGATCAGCAAAAGCAGG) as the primer^49^ and the virus RNA as the template. The cDNA was then used for qPCR (TaqMan Fast Advanced Master Mix, ThermoFisher Scientific) detailed previously^50^ by primers and probe targeting the NP segment which generate less internal deletion based on our previous study^50,51^. The relevant virus particle concentration (genome equivalent) was calculated based on the standard curve from the serial dilution of one cDNA from the virus.

### MUNANA

Virus was first diluted at least 5-fold in NA buffer (1% BSA + 33 mM MES + 4 mM CaCl_2_, pH = 6.5) to adjust pH and pre-warmed to 37 °C. 20 μL of diluted virus was added into a pre-warmed black 96-well plate with 20 μL of pre-warmed MUNANA (200 μM) in NA buffer. The V_mean_ of fluorescent signal (ex. 365nm, em. 450 nm) from 10 min to 45 min was obtained by the plate reader (Synergy HTX, BioTek) and used as the NA activity measured by MUNANA assay. The optimal range of reading was pre-determined. The NA activity was then normalized by the genome equivalent of the input virus.

### ELLA

The ELLA protocol was based on previous publication^52^. Briefly, 96-well plate was coated with fetuin. Virus was diluted in the NA buffer (same with MUNANA) added into the plate and incubated for 16-18 hours. Then, the plate was stained by horseradish peroxidase-conjugated peanut agglutinin (PNA-HRPO). The absorbent (A450) was measured by the plate reader after HRPO substrate (1-Step Ultra TMB-ELISA, ThermoFisher Scientific) was. NA activity was interpolated based on A450 and the standard curve from the serial dilution of a virus and normalized by the genome equivalent of the input virus.

### Deep mutational scanning of HA1

The generation, passaging and analysis procedure is based on our previous publication. Briefly, we first generated the plasmid library containing all the possible amino acid substitutions on HA1 subunit (D18-R344, H1 numbering from the first amino acid residue) by overlapping PCR (Phusion High-Fidelity DNA Polymerase, ThermoFisher Scientific) using primers containing the degenerative codon NNK. The fragments were then cloned into pDZ vector that express both the positive and negative sense of virus RNA. The HA1dms plasmid library, along with other seven segments, were co-transfected into HEK-293T cells co-cultured with MDCK-SIAT1. The supernatant was then collected 72 hours post transfection and used to infect MDCK-SIAT1 at MOI of 0.05.

The virus supernatant was harvested 36 hours post infection and prepared for barcode subamplicon sequencing. 140 μL virus supernatant was treated with 0.25 μg RNaseA for 30 min in 37 °C to remove the extracellular RNA and continued with viral RNA extraction (QIAamp Viral RNA Kits, Qiagen) following the manufacture’s protocol. The viral RNA product was then treated by DNase to remove the carry-over plasmid DNA (RNeasy Kit, Qiagen). cDNA of the virus genome was generated by reverse transcription with random hexamer as the primer (SuperScript III, ThermoFisher Scientific). The HA1 subunit was divided into three fragments for PCR (PrimeSTAR Max DNA Polymerase, Takara Bio), seven Ns and partial sequencing adapters were added in the primer as the barcode. The fragments were purified (PureLink Quick Gel Extraction Kit, invitrogen) and served as template for the second round PCR to add the full TruSeq adapters and purified (PureLink Quick Gel Extraction Kit, invitrogen) for NovaSeq. The libraries were pooled; quantitated by qPCR and sequenced on one SP lane for 251 cycles from both ends of the fragments on a NovaSeq 6000 with V1.5 sequencing kits.

The sequencing results were processed by dmstools2 using default parameters^53^. In the codon read sheets, codons with more than 20 reads in the WT plasmid were masked to reduce errors and then were translated into amino acid sheets. In the amino acid sheets, mutations with frequency less than 2×10^-5^ (∼20 reads) and less than 8 times more in the HA1dms plasmid library than the frequency in the WT plasmid were masked due to insufficient frequency for downstream analysis. The enrichment for each mutation is the fold change between the frequency post passaging and the frequency in the HA1dms library. The normalized relative fitness score was calculated by the equation below:

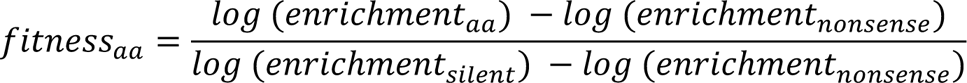

The receptor binding site is defined as follows: residues 133 to 138, 188 to 194, 221 to 228, 98, 153, 183 and 195^54^. The surface residues are identified based on solvent/protein contact surface by function “FindSurfaceResidues” on PyMOL.

### Antibody expression and purification

Plasmids encoding the heavy chain and light chain of the antibody were transfected at an equimolar ratio into Expi293F cells using ExpiFectamine 293 transfection kit (Gibco) according to the manufacturer’s protocol. Six days post-transfection, cell suspension was harvested, and the supernatant obtained through centrifugation at 4,500 × *g* and 4 °C for 45 min. The supernatant was clarified using a polyethersulfone membrane with a 0.22 μm filter (Millipore).

CH1-XL agarose bead slurry (Thermo Scientific) was washed with MilliQ H_2_O. Washed beads were added into the supernatant and incubated overnight at 4 °C with gentle rocking. Then, the supernatant containing unbound antibody was collected and discarded. Beads were washed with PBS. Subsequently, beads were incubated with 60 mM sodium acetate, pH 3.7 for 10 min at room temperature to elute antibody into 1 M Tris-HCl, pH 9.0. The antibody was buffer-exchanged into PBS using an Amicon ultracentrifuge filter with a 30 kDa molecular weight cutoff (Millipore) through four rounds of centrifugation at 3,000 × *g* and 4 °C for 10 min each. The antibody was further clarified using a Costar Spin-X centrifuge tube with a 0.22 μm cellulose acetate filter (Corning) through centrifugation at 10,000 × *g* and 4 °C for 10 min. Antibody concentration was measured with a Nanodrop spectrophotometer.

### Antibody selection

To determine the saturating neutralization concentration, 10^7^ TCID50 of CA09 was incubated with various concentrations of the antibody that was achievable for 30 min in 37 °C. The virus-antibody mixture was then used to infect the MDCK-SIAT1 cells in 6-well plates for 1 hour in 37 °C. The monolayer of cells was then washed twice with PBS and infectious medium supplemented with given concentration of antibody was added on top. The supernatant was then collected 24 hours post infection and titered by TCID50 assay. The saturating neutralization concentration was determined as the concentration where adding more antibody would not further decrease the yield of the virus or the highest concentration achievable.

In the selection experiment, the virus population was generated by mixing three independently grown virus stocks to ensure sufficient diversity in the starting population. Similar to the method used to determine the saturating neutralization concentration stated above, 10^7^ TCID50 of the recombinant viruses were neutralized by the antibody in the saturating neutralization concentration and used to infect the cells with 6 replicates. The virus was passaged to reach sufficient titer (> 10^4^ TCID50/mL) for next generation sequencing (NGS) if necessary.

To generate the library for NGS, the virus stock was first treated by RNase to remove the extra-cellular RNA and continued to viral RNA extraction (QIAamp Viral RNA Kits, Qiagen). 10 μL of viral RNA was added into 20 μL reverse transcription reaction using MBTuni-12 as the primer and high-fidelity reverse transcriptase (SuperScript III, ThermoFisher Scientific). The virus genome was further amplified by PCR (Phusion High-Fidelity DNA Polymerase, ThermoFisher Scientific) with 10 μL cDNA in each 50 μL reaction and MBTuni-12/13 (5′-ACGCGTGATCAGTAGAAACAAGG) as the primers for 25 cycles. The PCR product was then purified (PureLink Quick Gel Extraction Kit, invitrogen), prepared for NGS by shotgun library preparation (KAPA HyperPrep Kits, Roche), and then sequenced on either MiSeq (MiSeq 500-cycle sequencing kit version 2) or NovaSeq for 250nt-paired sequencing.

The sequencing results were first parsed by fastp (0.19.5) to remove low quality reads with a cutoff quality of 28 and minimum length of 100 nt^55^. After the quality filters, the reads were aligned to the reference sequence by bowtie2 (2.3.5.1)^56^ and transform to sorted bam file by SAMtools (1.9)^57^. The identity and the frequency of each mutation was identified by ivar (1.3.1) ^58^ using default parameters. To filter the major substitutions selected by the antibody pressure, only substitutions found twice in all the samples and reached 10% in frequency at least once were considered for further analysis.

## Data Availability

The raw sequencing data generated in this study was submitted to Sequence Read Archive (SRA) BioProject ID PRJNA1088522.

## ACKNOWLEDGMENTS

This study was generously supported by the National Institute of Allergy and Infectious Diseases of the National Institutes of Health under awards 1R01AI179910, 1R01AI139246, and sub-contract 75N93021C00017 through Emory-Center of Excellence in Influenza Research and Response to C.B.B.

We are grateful to Alvaro Hernandez, Chris Wright, and the DNA services team in Roy J. Carver Biotechnology Center for generating high quality sequencing results. We thank Dr. Patrick C. Wilson for generously providing the plasmids encoding some of the monoclonal antibodies used in this study. We thank Mireille Farjo and Lorenzo M. D’Alessio for helping us visualize the phylogenetic tree.

## APPENDICES

**Figure S1.**
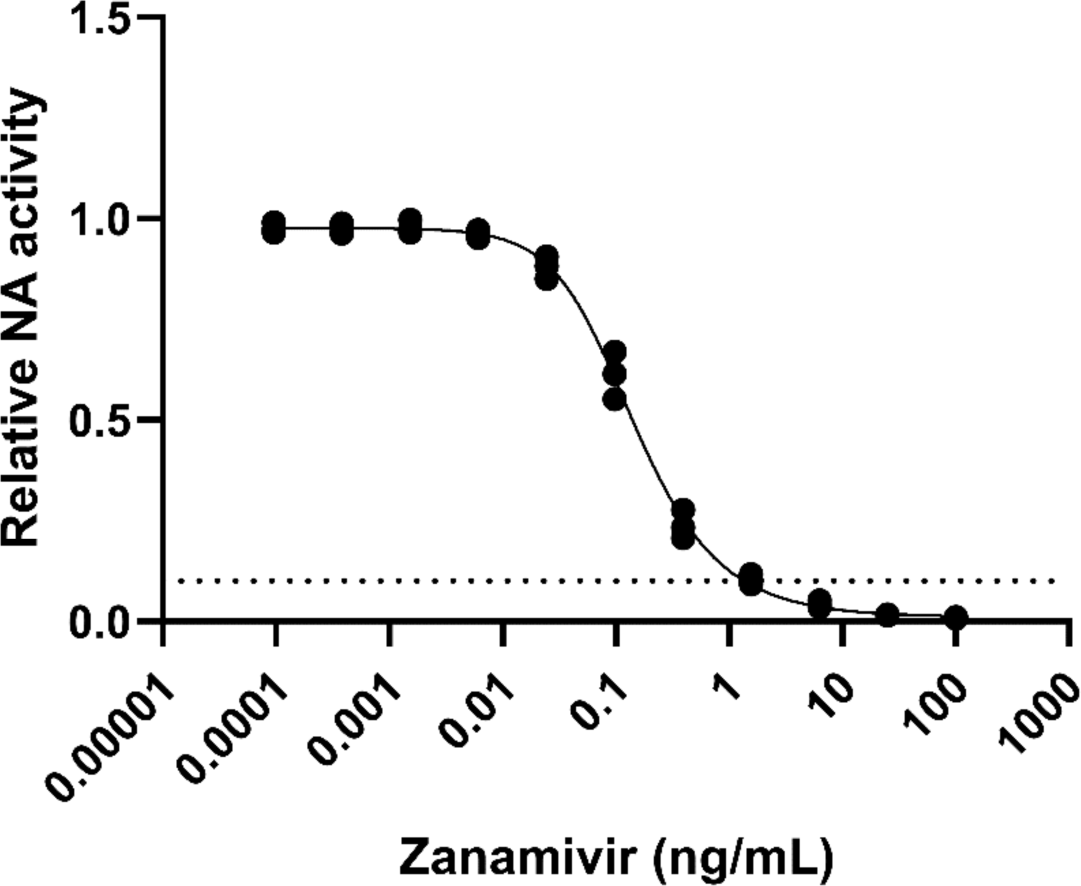
Inhibition curve of zanamivir against CA09 virus measured by MUNANA assay. Nonlinear curve fits by asymmetric sigmoidal in GraphPad Prism, dot line indicates 90% inhibition of NA activity, n = 3.

**Figure S2.**
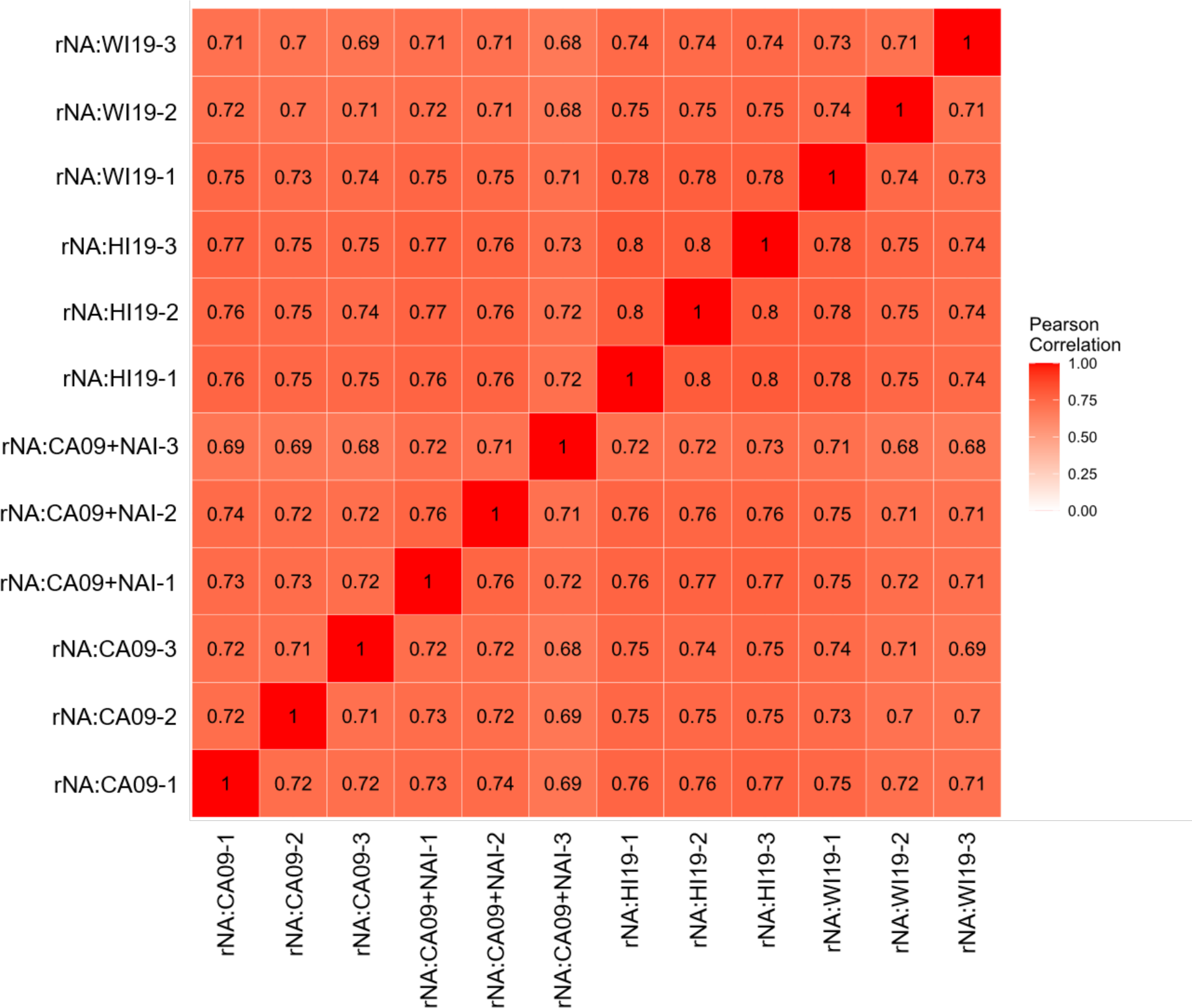
Pearson correlations of normalized relative fitness score between samples in deep mutational scanning.

**Figure S3.**
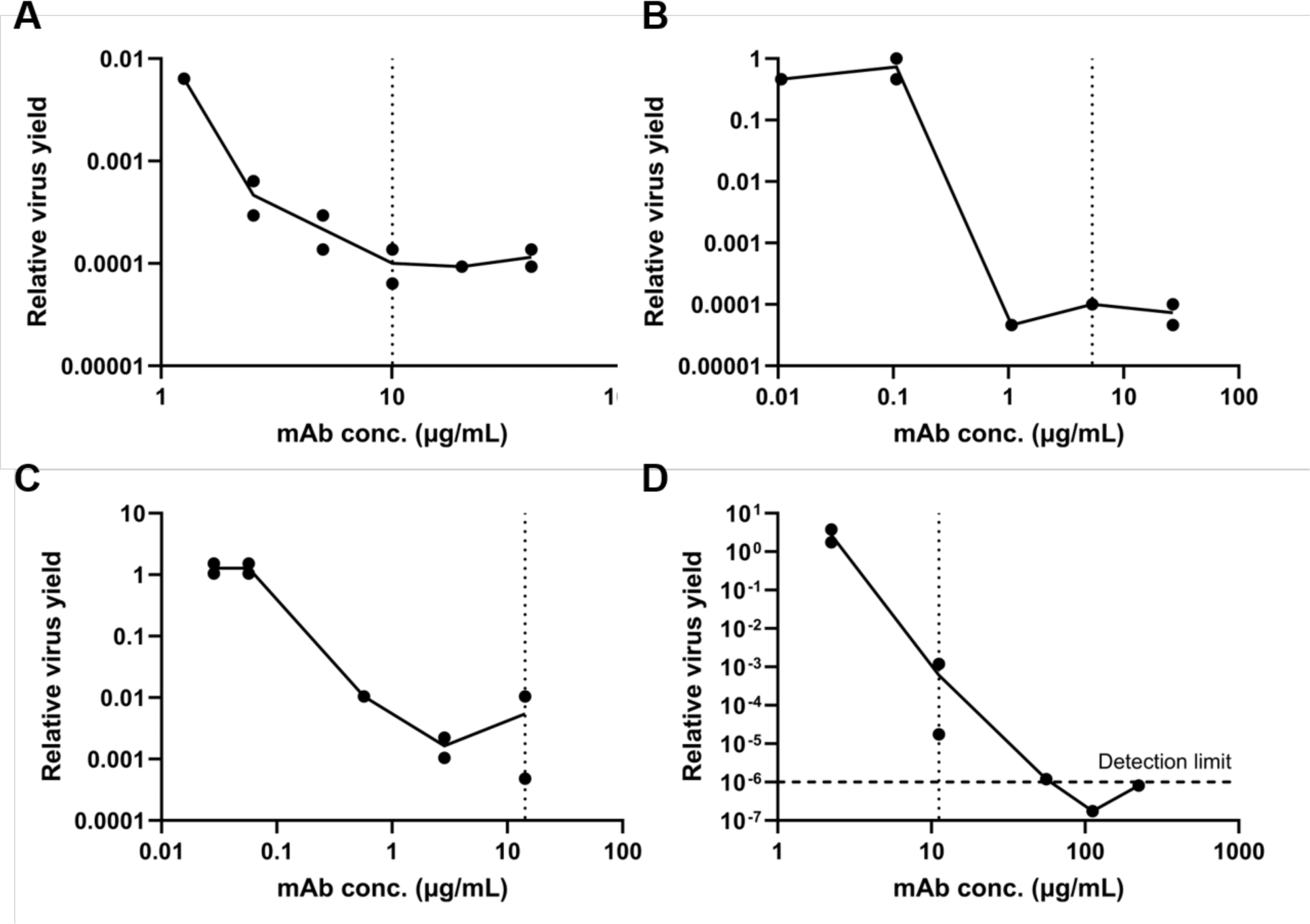
Saturated neutralization concentrations for the monoclonal antibodies used in this study. 10^7^ TCID50 of rNA:CA09 was neutralized with (A) EM4-C04, (B) 2A05, (C) 2C05 and (D) CR9114 in the given concentration and infected the cells. Virus supernatants were collected 24 hours post infection (n = 2) and measured by TCID50 assay. Relative virus yield was normalized to the no antibody control group. Dot lines indicate the concentration used in selection.

**Figure S4.**
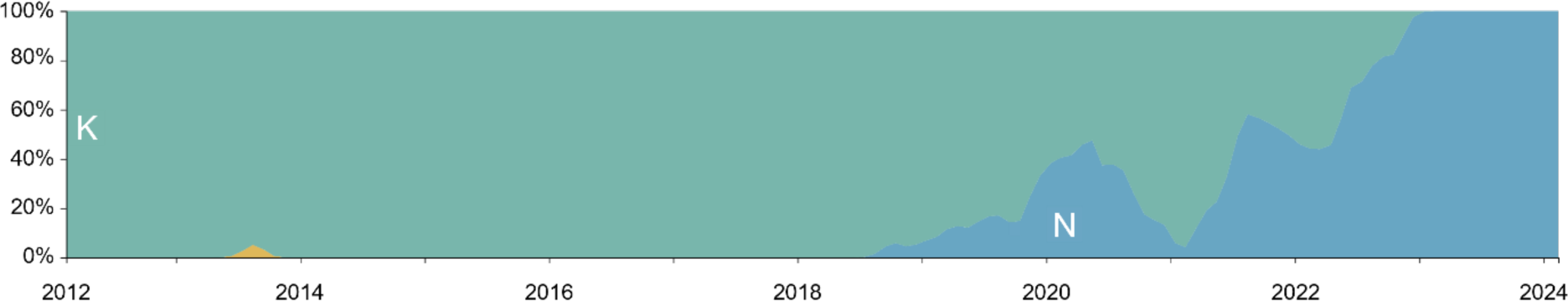
Normalized frequencies composition of pdmH1N1 HA position 133a over the years colored by the amino acid: lysine(K, green), asparagine (N, blue), arginine (R, yellow) from 1472 genomes samples between February 2012 to February 2024 (Screenshot taken from Nextstrain^34,59^).

**Figure S5.**
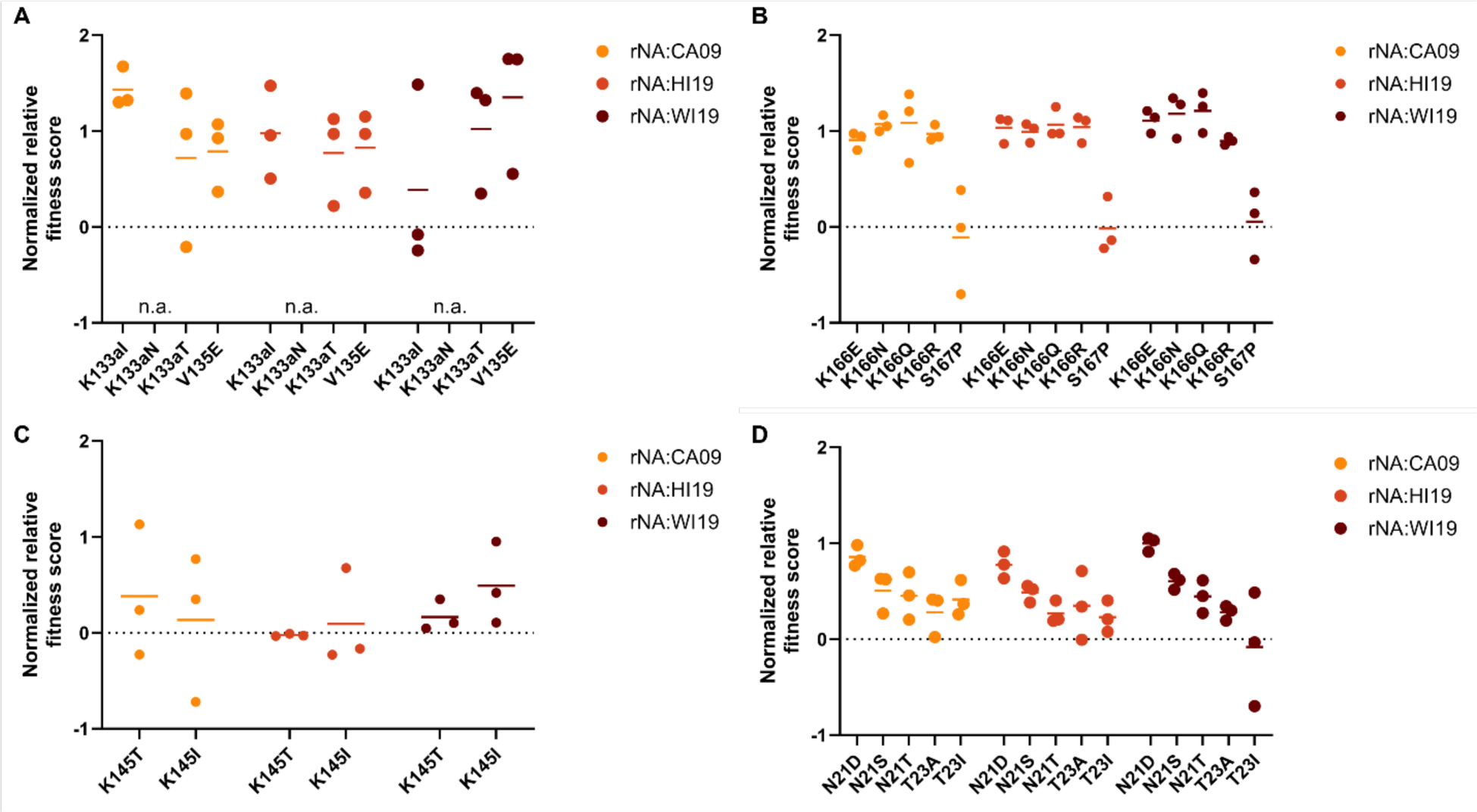
The normalized relative fitness score in deep mutational scanning of escape variants found in antibody selection with (A) EM4-C04, (B) 2A05, (C) 2C05 and (D) CR9114 (excluding HA2 residues).

